# Automated Long Axial Field of View PET Image Processing and Kinetic Modelling with the TurBO Toolbox

**DOI:** 10.1101/2025.06.06.658206

**Authors:** Jouni Tuisku, Santeri Palonen, Henri Kärpijoki, Aino Latva-Rasku, Nelli Tuomola, Harri Harju, Sergey V. Nesterov, Vesa Oikonen, Hidehiro Iida, Jarmo Teuho, Chunlei Han, Tomi Karjalainen, Anna K. Kirjavainen, Johan Rajader, Riku Klén, Pirjo Nuutila, Juhani Knuuti, Lauri Nummenmaa

## Abstract

Long axial field of view (LAFOV) PET imaging requires a high level of automation and standardization, as the large number of target tissues increases the manual workload significantly. We introduce an automated analysis pipeline (TurBO, Turku total-BOdy) for preprocessing and kinetic modelling of LAFOV [^15^O]H_2_O and [^18^F]FDG PET data, enabling efficient and reproducible analysis of tissue perfusion and metabolism at regional and voxel-levels. The approach employs automated processing including co-registration, motion correction, automated CT segmentation for region of interest (ROI) delineation, image-derived input determination, and region-specific kinetic modelling of PET data.

**Methods:** We validated the analysis pipeline using Biograph Vision Quadra (Siemens Healthineers) LAFOV PET/CT scans from 21 subjects scanned with [^15^O]H_2_O and 16 subjects scanned with [^18^F]FDG using six segmented CT-based ROIs (cortical brain gray matter, left iliopsoas muscle, right kidney cortex and medulla, pancreas, spleen and liver) representing different levels of blood flow and glucose metabolism.

**Results:** Model fits showed good quality with consistent parameter estimates at both regional and voxel-levels (R² > 0.83 for [^15^O]H_2_O, R² > 0.99 for [^18^F]FDG). Estimates from manual and automated input functions were in concordance (R² > 0.74 for [^15^O]H_2_O, and R² > 0.78 for [^18^F]FDG) with minimal bias (<4% for [^15^O]H_2_O and <10% for [^18^F]FDG). Manually and automatically (CT-based) extracted ROI level data showed strong agreement (R² > 0.82 for [^15^O]H_2_O and R² > 0.83 for [^18^F]FDG), while motion correction had little impact on parameter estimates (R² > 0.71 for [^15^O]H_2_O and R² > 0.78 for [^18^F]FDG) compared with uncorrected data.

**Conclusion:** Our automated analysis pipeline provides reliable and reproducible parameter estimates across different regions, with an approximate processing time of 1-1.5 h per subject. This pipeline completely automates LAFOV PET analysis, reducing manual effort and enabling reproducible studies of inter-organ blood flow and metabolism, including brain-body interactions.

## INTRODUCTION

Long axial field of view (LAFOV) PET systems present unique opportunities for scientific research and clinical diagnostics by enabling the simultaneous, noninvasive imaging of multiple organs and their physiological interactions. These systems provide clear benefits, including improved count sensitivity, extended image coverage, and extraction of image-derived input function from the aorta (*1*). However, they also introduce new challenges for data analysis. Patient movement is a major concern in LAFOV studies, as complete patient restraining is not possible similarly to e.g., in brain-only studies. In addition, manual pre-processing of LAFOV PET data is time-consuming, especially when analyzing multiple tissues. One LAFOV study may contain tens of different regions of interest (ROI), whose delineation alone can exceed a full workday per subject. Furthermore, the manual approach is prone to operator bias, which may reduce reproducibility compared to automated methods (*2*). Finally, the complex preprocessing, modelling and data analysis flow warrants comprehensive quality control to ensure accurate outputs at all stages and again, completing all these steps by human operators is simply not feasible.

Any all-inclusive LAFOV PET processing pipeline should handle preprocessing, ROI delineation, and kinetic modelling using various approaches tailored to different radiotracers, which makes the development of a comprehensive PET pipeline a challenging task. While various tools exist for automated brain PET data analysis (*2–5*), equivalent pipelines for LAFOV PET data are not yet widely available. To address these issues, we introduce an open-source Turku Total-Body PET modelling pipeline (TurBO), which enables automated and reproducible processing and kinetic modelling of LAFOV PET data. It supports various radiotracers and kinetic models, allowing fully automated processing, including co-registration, motion correction, input function determination, and ROI delineation. The pipeline enables both regional and voxel-level kinetic modelling, with the flexibility to apply tissue-specific models, and provides visual and numerical tools for quality control of the preprocessing steps.

Here, we describe and validate the TurBO pipeline (freely available at https://turbo.utu.fi) using [¹⁵O]H₂O and [¹⁸F]FDG PET data to benchmark preprocessing and modelling of tissue perfusion and glucose metabolism. We compared the automated image-derived input function (IDIF) with the manually delineated input, assessed results with and without motion correction, evaluated the results of ROI-based and voxel-level kinetic modelling, and compared outcome measures from manually and automatically determined ROIs from data collected with the Biograph Vision Quadra (Siemens Healthineers) 106 cm FOV PET/CT scanner. We hypothesized that TurBO provides a rapid, accurate and reproducible approach for modelling LAFOV PET perfusion and metabolism data at voxel and regional levels.

## MATERIALS AND METHODS

### Overview of TurBO pipeline

TurBO (Turku total-BOdy) pipeline runs on MATLAB (The MathWorks, Inc., Natick, MA, USA) and utilizes openly available tools for data processing. For all PET kinetic modelling, we use MATLAB implementations of openly available and previously validated in-house software (http://www.turkupetcentre.net/petanalysis). The general framework involves CT to PET registration, PET motion correction, segmentation of the CT image into tissues and organs, automatic image-based input function (IDIF) extraction and finally, kinetic modelling at regional and voxel-levels. The pipeline outputs LAFOV parametric images, separate parametric brain images normalized to MNI space, regional outcome measures as well as quality control metrics. An overview of the workflow is shown in **Figure 1**, and the process is described in detail below.

**Figure 1.**
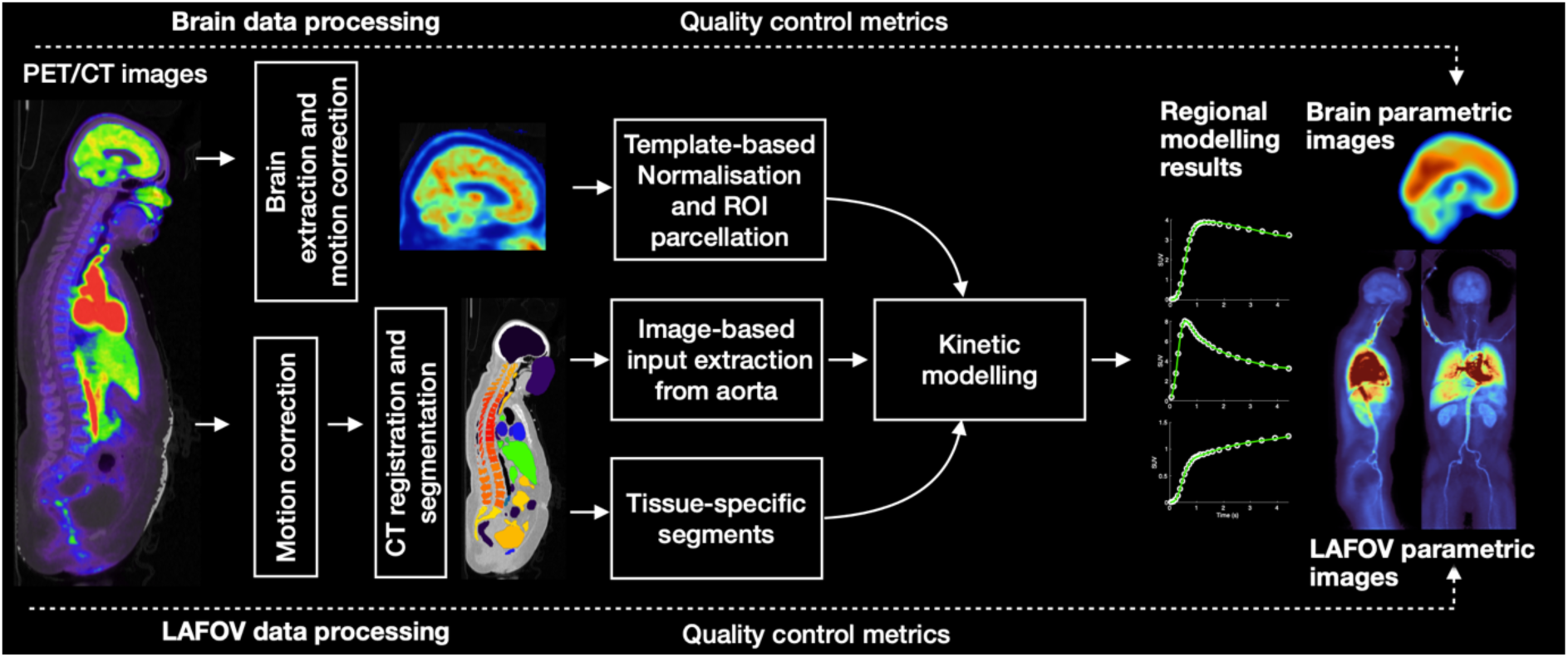
Flowchart for the TurBO pipeline.

### LAFOV PET data preprocessing and kinetic modelling

DICOM images are first converted to NIfTI using the *dcm2niix* tool (*6*), and the required metadata such as framing, injected dose, and subject weight is extracted from the DICOM header. Next, PET motion correction is applied following the previously described approach (*7*), employing a diffeomorphic *greedy* registration algorithm (https://greedy.readthedocs.io). For [^18^F]FDG, the motion correction start frame is determined by normalized cross-correlation (NCC), as described in (*7*). In our [^18^F]FDG validation data this varies between 4 - 6 min. Due to the variation in [^15^O]H_2_O distribution, early frames are discarded, and the start frame is selected as the time point where the heart time-activity curve (TAC) falls below half of its peak value (this varies between 15 – 45 s in our validation data). For both radioligands, the motion correction reference frame is selected as the one with the highest NCC relative to the CT. In our [^15^O]H_2_O validation data, this frame is typically near the scan midpoint and in [^18^F]FDG data near the scan endpoint. Subsequently, CT is registered with the PET mean image using the greedy algorithm to correct any misalignments and ensuring that CT-based regions of interest (ROIs) correspond with the PET data. The CT image is resampled to PET voxel size, whereafter the *TotalSegmentator* tool (*8*) is used for CT-based segmentation of major organs and tissues, to serve as ROIs for PET quantification. Kinetic modelling for [^15^O]H_2_O is carried out using one tissue compartment model (1TCM), while [¹⁸F]FDG analysis includes Patlak plot (*9*), fractional uptake ratio (FUR) (*10*) and standardised uptake value (SUV) methods (see **Supplementary material** for full details).

#### Image-derived input extraction

For automatic IDIF determination, a separate smaller PET image containing the heart and aorta is extracted from the original LAFOV PET image and corrected for motion using the rigid motion correction method (*11*), which enables motion correction also for the initial frames. Consequently, IDIF is extracted from the descending aorta using an automated method based on our center’s standard protocol for manual input delineation: the lower third of the myocardium is used as a higher anatomical landmark to identify the starting point. From this level, the maximums are located from each axial slice and connected transaxially by gap-filling algorithm 10 cm downwards to form a continuous, string-like ROI along the aorta. (**Figure 1 A-B**).

#### Brain-PET data analysis

The brain consists of distinct cytoarchitectonical and functionally separable regions, which are previously defined in various atlases in standard stereotactic space. Therefore, a separate analysis pathway is used to incorporate these atlases and to facilitate whole-brain analysis using widely used toolboxes (e.g., SPM, FSL) based on parametric mapping of spatially normalized brain images. In this stream, a freely available in-house Magia-toolbox (*2*) is used for extracting the brain from the original PET image, followed by rigid motion correction and spatial radioligand template-based normalization for transforming the desired atlas from standard MNI space to the subject native space. The pipeline includes the ROIs from AAL-atlas (*12*) containing cortical lobes and selected subcortical regions, but any other atlas in MNI space can also be used. Finally, kinetic modelling (see **Supplementary material** for details) is carried out for each region, as well as at the voxel-level for the whole brain data.

#### Output

As a final step in the processing, the parameter estimates, CT-based ROI volumes, and model goodness of fit metrics (Pearson’s R^2^) are saved for each region in the standard format. These results can then be retrieved for selected subjects using a dedicated function. Regional results from voxel-level parameter maps and from clustered regions can also be collected similarly.

The voxel-level parameter maps are also clustered into three subregions within selected areas using hierarchical clustering. This supports regional analysis of functionally distinct areas, such as the kidney cortex and medulla or brain grey and white matter. Additionally, the CT-based ROIs can be trimmed by removing a specified distance from their edges to reduce signal contamination from nearby high-intensity areas, or due to motion. For instance, it may be necessary to exclude spill-in from the cardiovascular system in the segmented liver.

Finally, quality control plots illustrating the CT to PET registration, corrected PET motion, regional model fits and overlay-images of CT and voxel-level parameters are saved in an html-file for later visual check (see **Supplementary material** for example of quality control output).

### Validation

We validated the automated [¹⁵O]H₂O and [¹⁸F]FDG analysis pipeline by comparing automatically and manually derived input functions, assessing the agreement between regional and voxel-level modelling, evaluating the effect of motion correction, and comparing the results from manual and automatic ROI delineation.

For validation purposes in the main report, we chose six segmented CT-based ROIs (cortical brain gray matter (GMctx), left iliopsoas muscle, right kidney cortex and medulla, pancreas, spleen and liver) representing different levels of perfusion and glucose metabolism. The results from all other segmented regions (n=72; ribs and vertebrae are excluded for clarity), are shown in the **Supplementary material**. To account for the contribution of arterial blood, perfusion was quantified as H₂O_flow_ = K_1_(1-V_A_), and glucose metabolism was quantified as Patlak K_i_. Because the dual input [^15^O]H_2_O model in the liver may have parameter identifiability issues in K_1_ and k_2_ parameters, we replicated the quantification using distribution volume K_1_/k_2_ (see **Supplementary material** for results).

The accuracy of the automatically derived descending aorta IDIF was tested by comparing the model parameter estimates from segmented CT-based ROIs using both automated and manually drawn input, which was delineated using the above-described criteria with Carimas software (*13*). Agreement between regional and voxel-level estimates was evaluated by comparing parameters estimated from averaged regional TACs to voxel-level estimates, which were averaged within each ROI. To assess the motion correction, parameter estimates from motion-corrected and uncorrected data were compared using CT-based ROIs. Finally, using descending aorta IDIF, the modelling results from three manually drawn axial ROIs (in liver, kidney cortices, and spleen) for [^15^O]H_2_O and two ROIs (iliopsoas muscle and liver) for [^18^F]FDG were compared with the corresponding results from automatically segmented CT-based ROIs.

Because the manually drawn ROIs were substantially smaller than the automatically segmented ones, we also compared the voxel-level results using manual ROI in the spleen and liver with results from CT-based ROIs where the volume was reduced (20 mm from the ROI borders in the liver and 10 mm in the spleen). Similarly, in the kidneys, the manual ROI results were compared to those from a functionally distinct PET-based cluster in the kidney cortex, and in the brain, PET-based cortical GMctx cluster results were compared with results of the corresponding AAL-atlas ROI from the PET-template-based brain processing. For [^15^O]H_2_O, we also compared myocardial blood flow (MBF) measured as k_2_ with descending aorta IDIF to manually assessed MBF using Carimas software with left ventricle input (*14*).

All regional [^15^O]H_2_O parameter estimates were calculated using non-linear least squares (NNLS) estimation with 100 randomly initialized parameters from the following lower and upper bounds: K_1_:[0, 1800] ml/(min*dl), K_1_/k_2_:[0, 1], V_A_:[0, 0.8]. Voxel-level [^15^O]H_2_O parameters were estimated using 500 basis functions with uniformly distributed k₂ values from the range [0, 6] 1/min.

### Validation Data

The automated processing and kinetic modelling for perfusion imaging was validated using LAFOV [^15^O]H_2_O PET data from 21 healthy subjects, which were acquired at Turku PET centre with Biograph Vision Quadra (Siemens Healthineers) PET/CT scanner with spatial resolution of 3.3-3.8 mm FWHM and an axial field of view of 106 cm (*15*). Validation for imaging the glucose metabolism was carried out using the same scanner with [^18^F]FDG PET data from healthy control and obese subjects (n=16). Data acquisition and subject’s demographic details and are described in the **Supplementary material**. The study was approved by the institutional ethical review board and conducted following the principles of the Declaration of Helsinki. All participants provided written informed consent prior to the examinations.

### Statistical methods

All statistical comparisons were carried out in R (version 4.4.1) using Pearson’s correlation and Bland-Altman analysis describing the relative difference and 95% limits of agreement between the methods. Paired t-test was used to assess the significant differences between the methods, where p<0.05 indicated a statistical significance.

## RESULTS

### Processing time

The typical total processing time for one subject was 61 min for [^15^O]H_2_O data, and 85.5 min for [^18^F]FDG data (**Supplementary material, Table S2)**.

### Model fits

Visual inspection indicated good model fits in in all regions, except in kidneys, where the model underestimated the measured TAC in the later time points. There was a moderate variability between subjects, but data and model fits had a high correlation across subjects (mean R^2^ > 0.88 for [^15^O]H_2_O, and mean R^2^>0.92 for [^18^F]FDG) in all studied regions (**Figure 3**).

### Automated versus manual ROI delineation

There was moderate overlap between the manually drawn and automatically derived input function mask images for both datasets, and in [^18^F]FDG data the manual input volume was larger compared to IDIF (**Supplementary material, Table S3**). Despite this, the areas under the curves (AUCs) of manually derived input function and IDIF (**Figure 2 C-D**) were almost perfectly correlated (R^2^=0.99).

**Figure 2.**
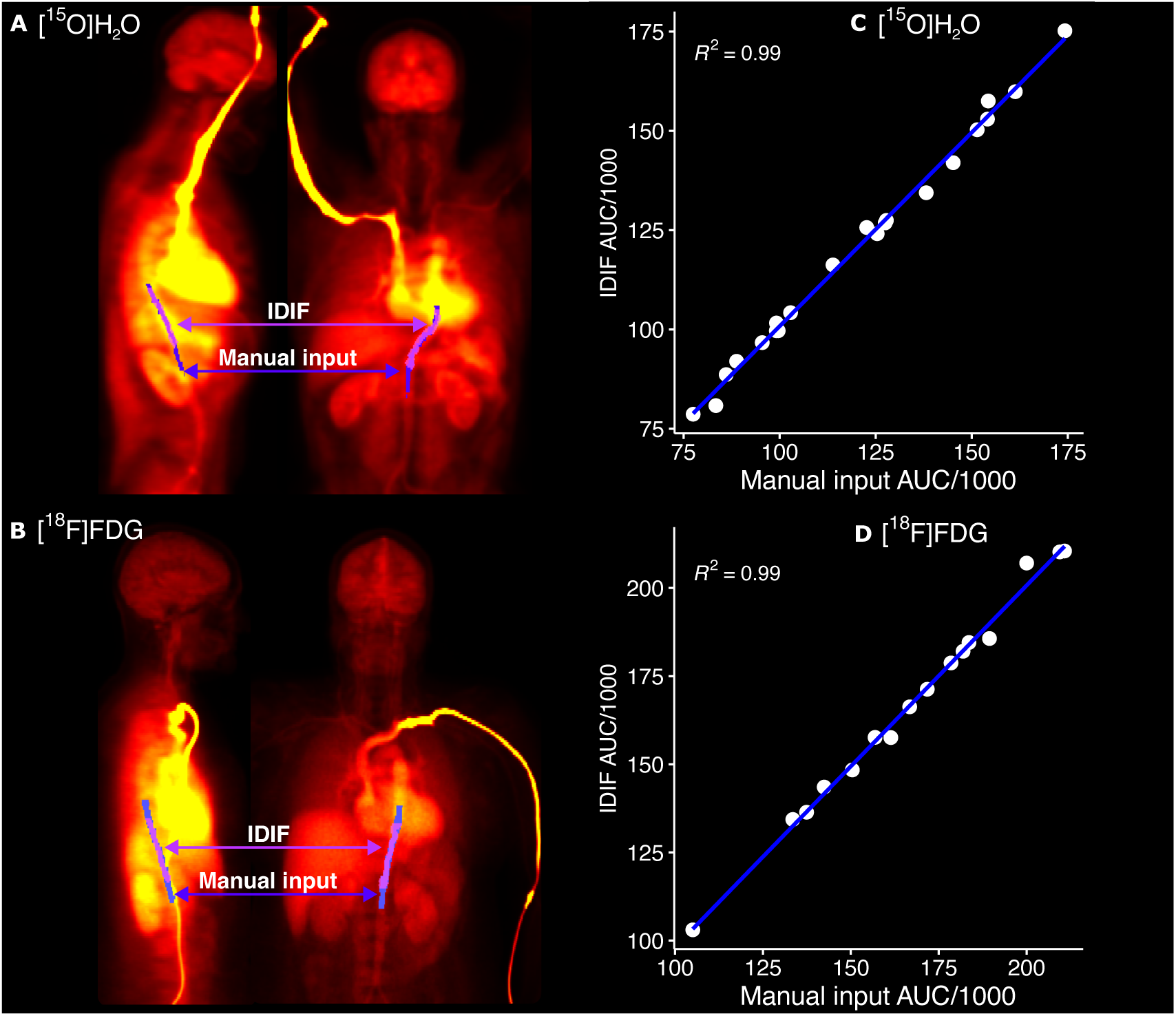
**A-B)** Manually delineated descending aorta ROI (blue) overlaid with image-derived input (violet) for [^15^O]H_2_O and for [^18^F]FDG. **C-D)** Scatterplots of areas under the input curves of manually delineated descending aorta and image-derived input for [^15^O]H_2_O and for [^18^F]FDG.

**Figure 3.**
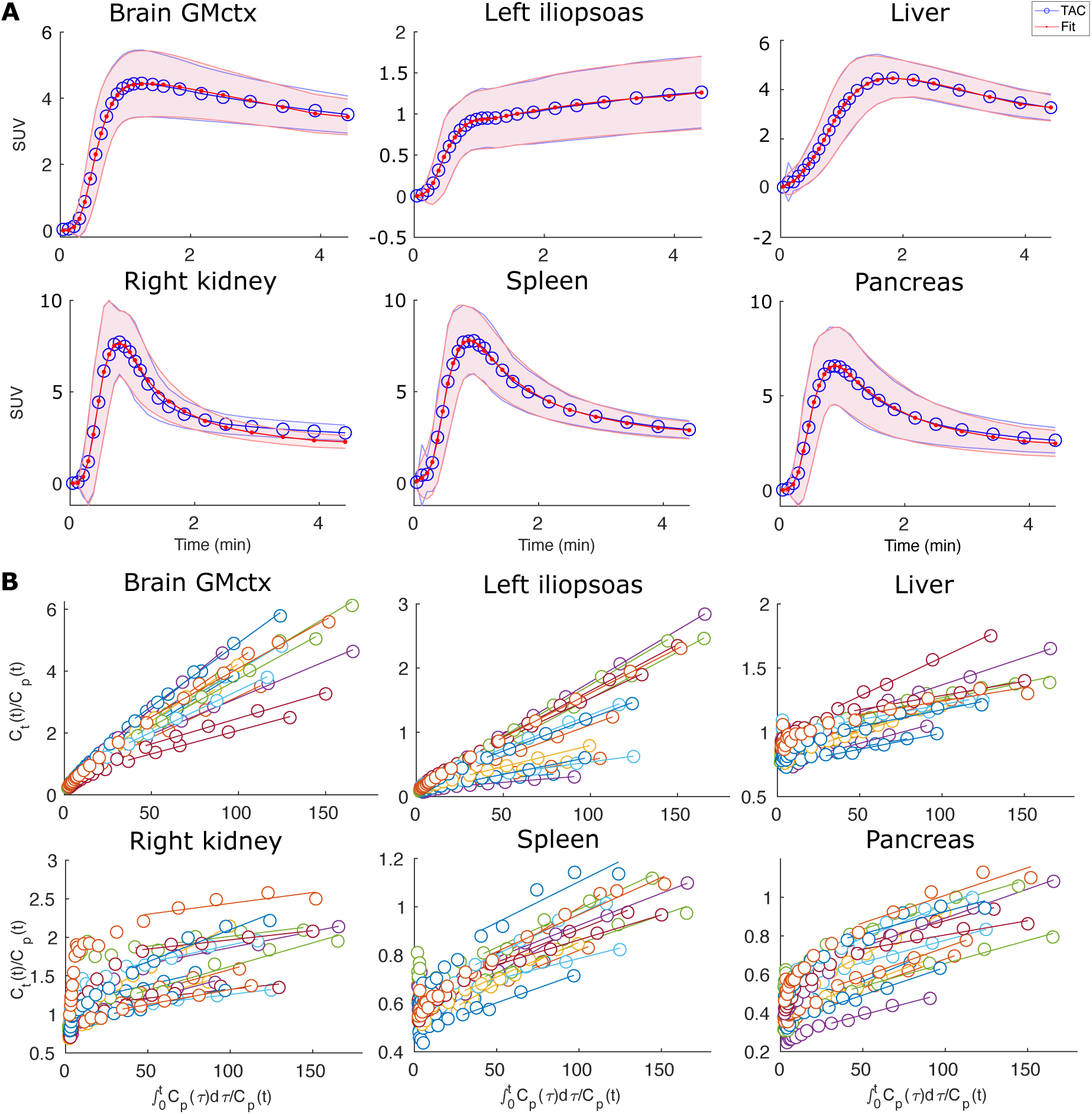
**A)** Between subject mean [^15^O]H_2_O PET time-activity curves (blue) and the mean one tissue compartment model fits (red) in brain cortical gray matter (GMctx), left iliopsoas muscle, liver, right kidney, spleen and pancreas for 21 subjects scanned at rest. Shaded areas illustrate the standard deviation of the data and the corresponding model fits. **B)** [^18^F]FDG Patlak-plots in GMctx, left iliopsoas muscle, liver, right kidney, spleen and pancreas for 16 subjects (shown in different colors).

The parameter estimates using manual and automatically derived input functions had high correlation (R^2^>0.74 for [^15^O]H_2_O, and R^2^>0.78 for [^18^F]FDG) in all selected six regions, with negligible bias for both radioligands (**Figure 4: manual vs. automatic input, Supplementary material, Figure S4**). When all regions were considered, the correlation and bias results were similar, but lower correlation and higher mean relative difference were observed in arteries and in heart and lung subregions mainly for [^15^O]H_2_O **(Supplementary material, Figure S5).**

**Figure 4.**
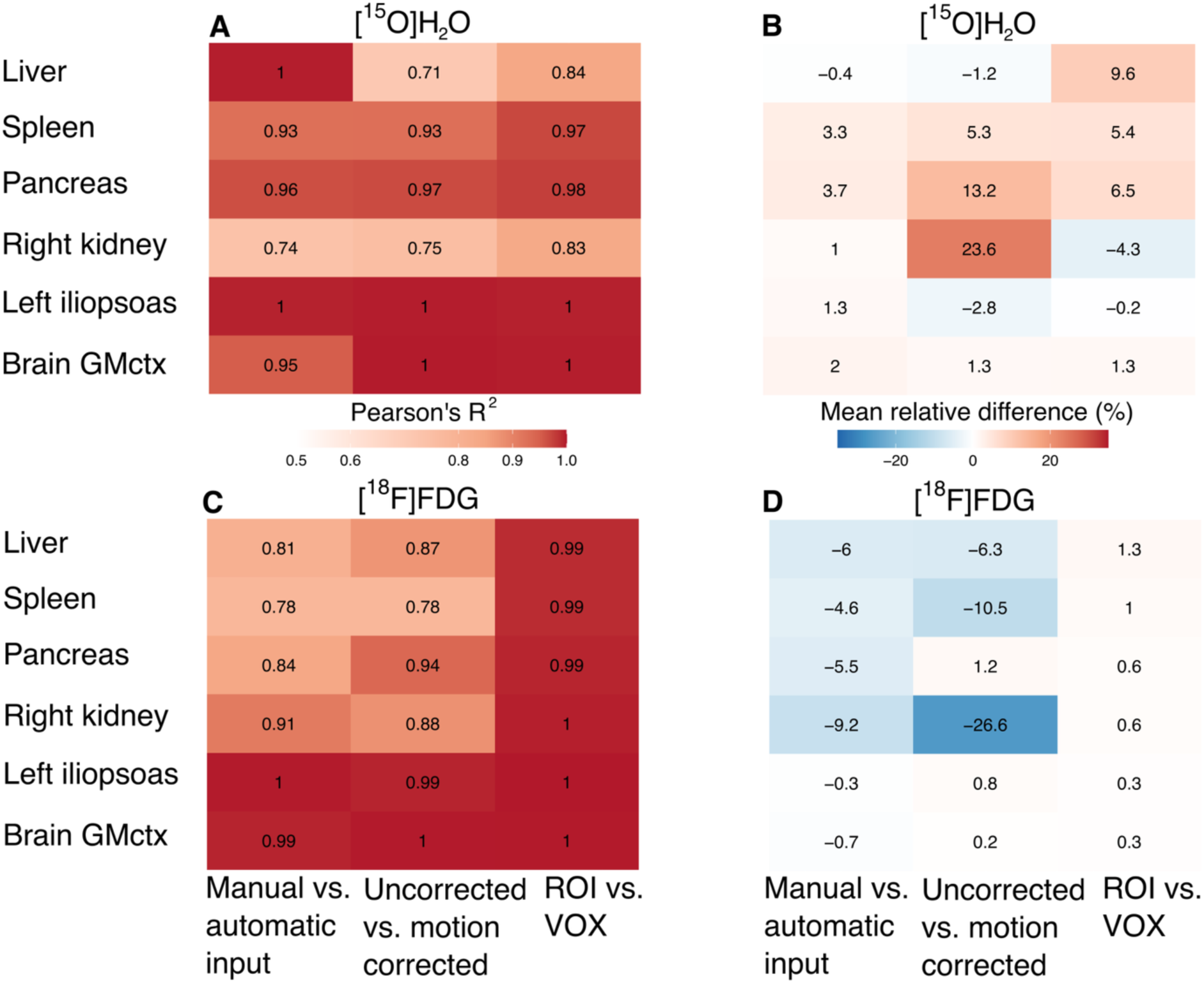
Comparisons between manual vs. automatic input determination methods, motion corrected and uncorrected data, and regional-level versus voxel-level (ROI vs. VOX) methods. **A)** Correlation coefficients (Pearson’s R^2^) and **B)** mean relative regional differences (%) of regional [^15^O]H_2_O parameter estimates. **C)** Correlation coefficients (Pearson’s R^2^) and **D)** mean relative regional differences (%) of regional [^18^F]FDG Patlak K_i_.

### Motion correction

To estimate the regional motion for each subject, we tracked the Euclidean distance of a single voxel, located at the mean of x-, and the maximum of y-, and z-coordinates in each segmented ROI, between the data with and without motion correction, and the results of corrected frame-wise motion for each ROI are summarized in **Supplementary material, Table S4**. Despite of the motion, parameter estimates using motion-corrected data were associated (R^2^>0.71 for [^15^O]H_2_O, and R^2^>0.78 for [^18^F]FDG) with the estimates obtained using uncorrected data (**Figure 4: uncorrected vs. motion corrected**). The mean relative difference varied by region, with higher difference in the right kidney (24%) and in pancreas (13%) in [^15^O]H_2_O data and in the right kidney (−27%) and in spleen (−11%) in [^18^F]FDG data, compared to the other selected regions. When all regions were considered, lower correlation and higher mean relative difference were found mainly in arteries and in the heart **(Supplementary material, Figure S6).**

### Regional versus voxel-level results

Despite significant differences in brain for both [^15^O]H_2_O and [^18^F]FDG, and in pancreas and spleen for [^15^O]H_2_O, the regional and voxel-level modelling results were highly correlated (R^2^>0.83 for [^15^O]H_2_O, and R^2^>0.99 for [^18^F]FDG) for both radioligands (**Figure 4: ROI vs. VOX, Figure 5)**. The mean relative difference between regional and voxel-level [^18^F]FDG modelling was less than 2%, and below 10% for [^15^O]H_2_O. When all regions were considered, lower correlation and higher mean relative difference were observed only in [^15^O]H_2_O data in arteries, lungs and in heart **(Supplementary material, Figure S7).**

**Figure 5.**
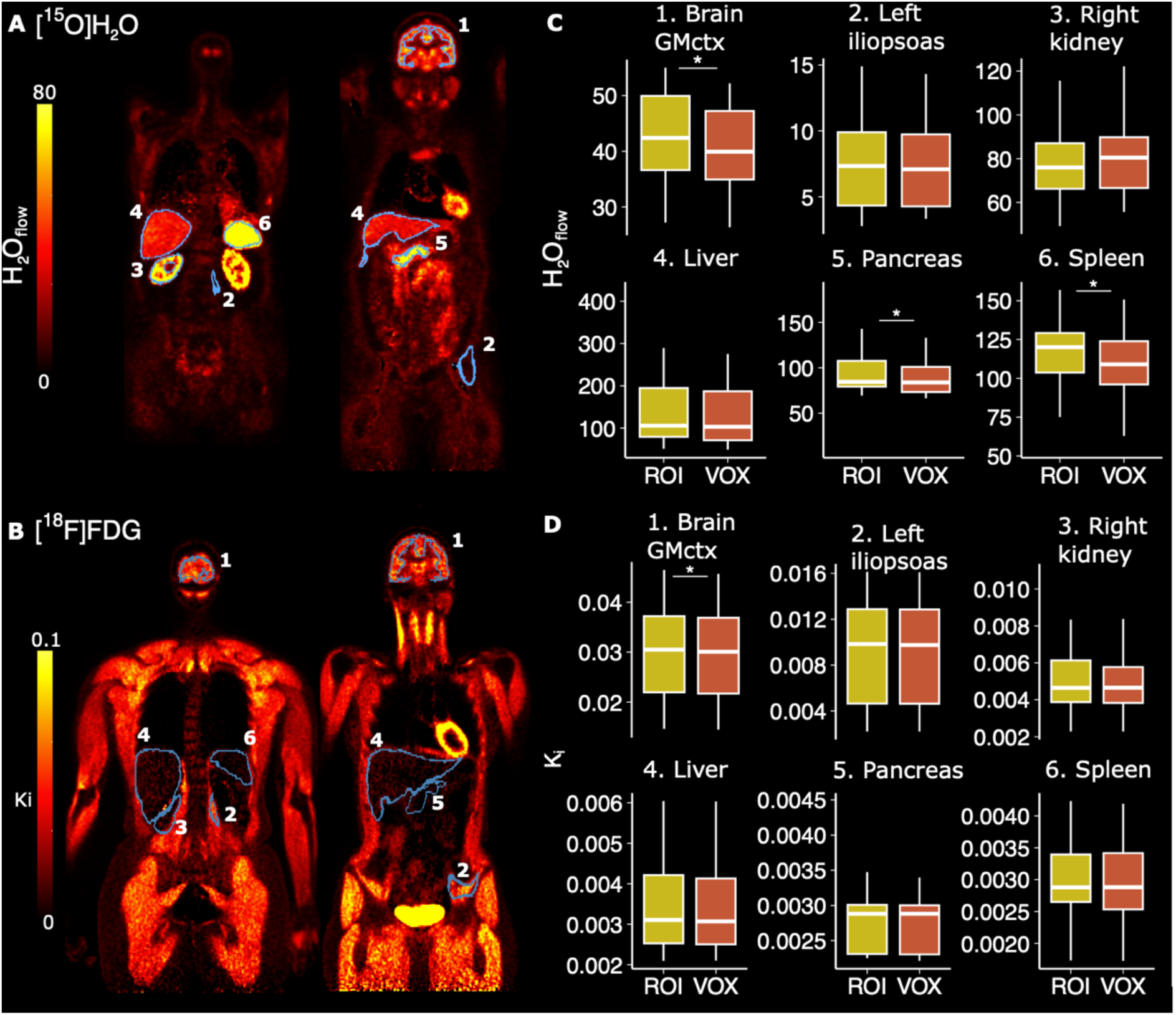
Voxel-level average intensity projection images for representative subjects: **A)** H₂O_flow_ image for [^15^O]H_2_O and **B)** K_i_ image for [^18^F]FDG. **C-D)** Boxplots illustrating the regional (ROI) and voxel-level (VOX) estimates in brain cortical gray matter (GMctx), left iliopsoas muscle, right kidney, pancreas and spleen for [^15^O]H_2_O (n=21) and [^18^F]FDG (n=16) data. Significant differences based on paired t-test are indicated with asterisks.

Voxel-level results obtained using the basis function method yielded higher correlation, and less bias with the ROI-level modelling results compared to the results of the voxel-level model, where all three parameters were estimated (Supplementary material, Figure S8).

Liver distribution volumes (K_1_/k_2_) showed similar results in all validation tests 1-3, as compared with the H₂O_flow_ (K_1_(1-V_A_)) estimates (Supplementary material, Figure S9), but with a significantly lower coefficient of variation (6.8% in K_1_/k_2_ vs. 65.8% in H₂O_flow_ estimates).

Manually drawn ROIs were substantially smaller than the automatically segmented CT-based ROIs (Supplementary material, Table S5). However, comparison between [^15^O]H_2_O and [^18^F]FDG parameter estimates from manually delineated ROIs and segmented CT ROIs showed high correlation (R^2^>0.82 for [^15^O]H_2_O, and R^2^>0.83 for [^18^F]FDG), but moderate mean relative differences (18% in kidney, 21% in spleen, 10% in liver and 8% in myocardium for [^15^O]H_2_O, and -19% in liver and 7% in iliopsoas muscle for [^18^F]FDG; **Figure 6**).

**Figure 6.**
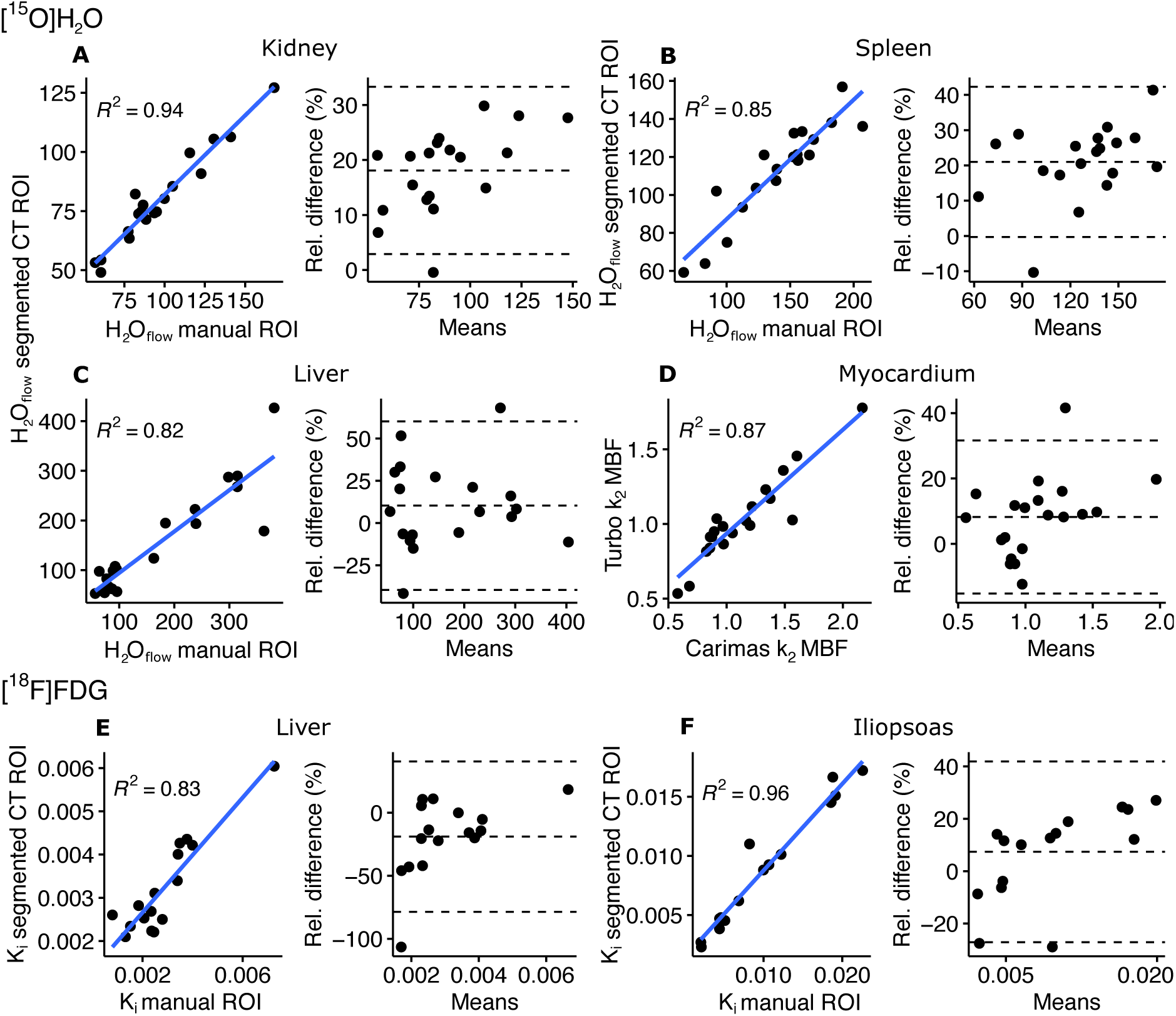
**A-C)** Scatterplots and Bland-Altman plots illustrating the correlation and relative difference between [^15^O]H_2_O parameter estimates of interest in manually drawn ROIs and segmented CT-based ROIs. **D)** Scatterplot and Bland-Altman plot illustrating the correlation and relative difference between myocardial blood flow (MBF) measured as k_2_ with descending aorta IDIF and manually assessed k_2_ MBF with left ventricle input. **E-F)** Scatterplots and Bland-Altman plots illustrating the correlation and relative difference between [^18^F]FDG Patlak K_i_ in manually drawn ROIs and segmented CT ROIs.

After reducing the volumes of segmented liver and spleen CT ROIs, the voxel level parameter estimates were more closely aligned with the results from manually drawn ROIs in the liver for [^15^O]H_2_O, but there were significant differences in the spleen for [^15^O]H_2_O and in liver for [^18^F]FDG based on paired t-test (**Figure 7**). Similarly, PET-based clustering gave corresponding estimates in the brain as compared to the results of PET-template based processing, but in the kidney, the clustered cortex ROI produced significantly higher estimates compared to the manually drawn cortex ROI for [^15^O]H_2_O (**Figure 7**).

**Figure 7.**
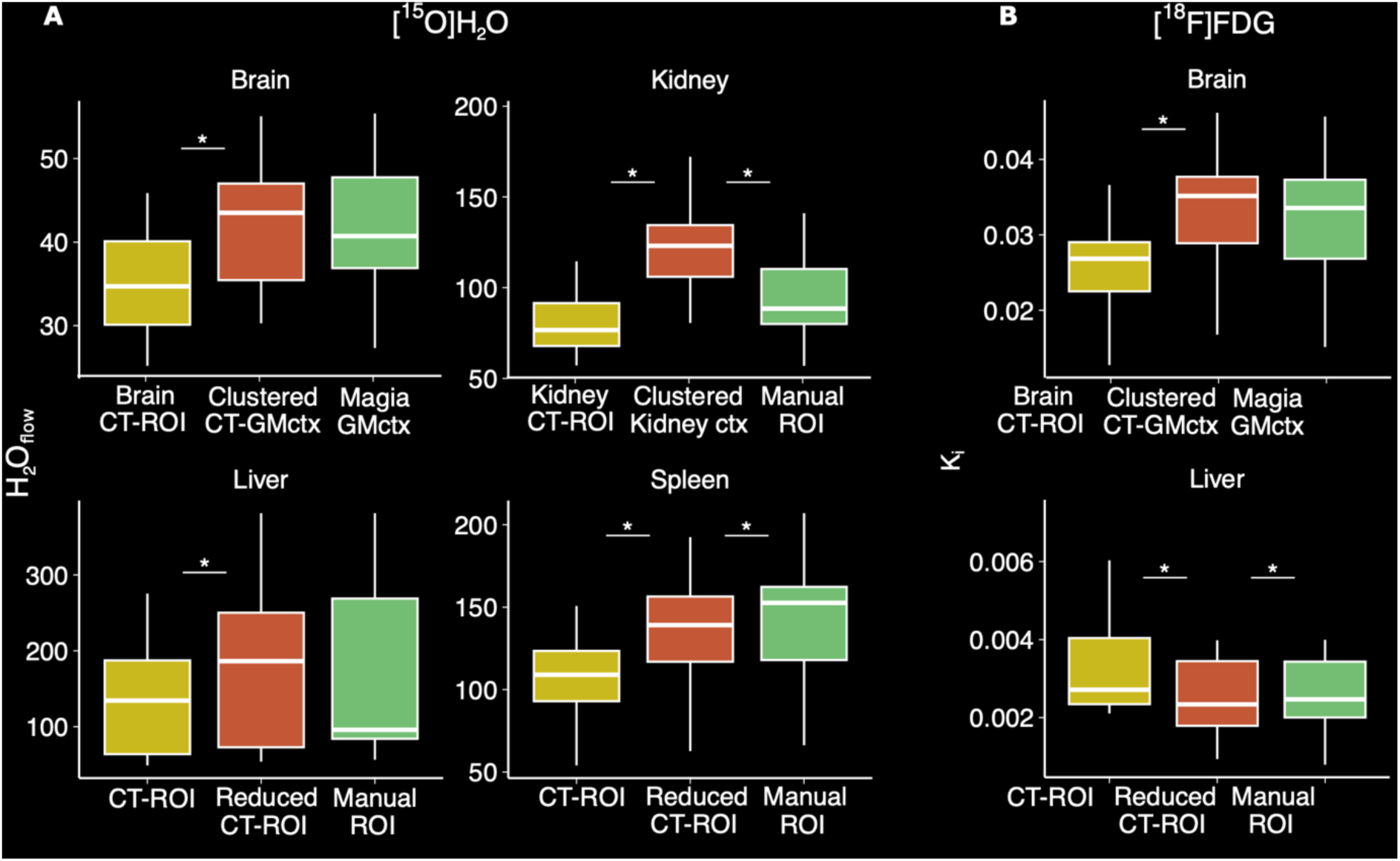
Boxplots illustrating **A)** [^15^O]H_2_O (n=21) and **B)** [^18^F]FDG (n=16) voxel-level results using manually drawn ROIs, CT-based ROIs, CT-based ROIs where the volume was reduced (in liver and spleen), and PET-based clustered ROIs (in brain and kidneys). Significant differences based on paired t-test with respect to clustered- and reduced ROIs are indicated with asterisks.

## DISCUSSION

We developed a unified pipeline for LAFOV PET data processing and kinetic modelling and demonstrated that it produces consistent regional estimates of radiotracer uptake for both tested radioligands [^15^O]H_2_O and [^18^F]FDG. In contrast with existing pipelines that are limited to specific tissues such as the brain or can handle only a single step of the data preprocessing such as kinetic modelling (*16*,*17*), our pipeline automates the full LAFOV workflow from pre-processing (image registration, segmentation, and motion correction) to kinetic modelling, making it more versatile and comprehensive compared to the other pipelines (*2*,*4*,*5*). This complete automation of the LAFOV PET data preprocessing and voxel- and region-level kinetic modelling provides significant advantages for the data analysis by improving the reproducibility of the analysis through automation and logging of the processing steps and parameters. It also enables efficient region-wise studies by removing the operator load through automation and by reducing inter-operator variability in ROI delineation. Overall, this approach allows standardized large-scale analysis of LAFOV PET data, paving the way for harmonized analysis and data integration in multi-center studies.

### Robust automated input delineation and modelling in regional and voxel-level

PET data kinetic modelling requires the input function typically obtained either from blood samples or from the image. Blood sampling introduces additional labour to the imaging protocol, but with sufficiently long axial FOV and short imaging frames, the input can be derived reliably from the aorta if it is visible in the image (*18*). Our results confirm that the automatic input delineation implemented in the TurBO pipeline is robust and comparable with manual delineation, and thus it can be reliably used instead of time-consuming manual input ROI drawing.

We also established that the regional outcome measures had high correlation between manually and automatically derived ROIs, and between voxel-level and region-level analyses. Voxel-level [^15^O]H_2_O modelling with basis functions improved alignment with the ROI-based results, when compared to the full three-parameter voxel-level model. However, this method requires more processing time because the delay parameter must be estimated for each voxel. Nevertheless, all processing and modelling for one subject was carried out in an hour for [^15^O]H_2_O and in less than 1.5 hours for [^18^F]FDG. With parallel computing, larger batches can be completed in a day, depending on the available resources.

Altogether, these results show, that TurBO provides a reproducible, reliable and computationally fast approach for modelling and investigating LAFOV PET data and facilitates the investigation of inter-organ interactions in tissue perfusion and metabolism.

### Motion correction

While controlling the subject motion is crucial to obtaining high-quality data, involuntary motion, such as breathing and cardiac motion is still present during image acquisition, as observed in our motion correction results. Although the motion during PET scanning and misalignment between CT and PET should ideally be corrected during image reconstruction, our pipeline also includes post-reconstruction correction methods. To account for complex body movements, we included in our pipeline a diffeomorphic correction method (*7*), that employs both rigid and nonlinear deformations. Our findings showed a high correlation between corrected and uncorrected data, but noticeable differences in absolute values, especially in the kidney and pancreas, which highlights the importance of motion correction.

Although previous research suggests that cardiac motion may require nonlinear correction methods (*19*), the recent findings have showed that rigid correction methods may be sufficient (*20*). For practical reasons, we used only rigid motion correction for the heart and neighbouring aorta, to minimize altering the original data, and to avoid possible image distortions from diffeomorphic correction. Based on our results the automated IDIF produced comparable results with the manually derived input without motion correction. The remaining small bias between the results of the automated and manual methods is likely caused by differences in the input ROI volumes, and because the subject motion is corrected in the automated IDIF. Apart from the input, the other manually drawn ROIs were substantially smaller than the automatically segmented CT-based ROIs, because the manual ROIs were drawn only to a limited number of slices. This likely explains the relative differences between the regional results from automatically derived and manual ROIs. Because the CT-based segmented ROIs are large entities, the pipeline includes also additional reduced ROIs, which mitigate the effects of subject motion and spill in from other nearby regions. Also, hierarchical clustering of voxel-based parametric maps within a selected region into separate subregions allow to examine the functionally distinct subregions, such as the kidney cortex and medulla.

### Limitations

The radiowater model in the liver may have issues with parameter identifiability due to the dual input model (*21*). Although the validation results were similar between H₂O_flow_ (K_1_(1-V_A_)) and distribution volume K_1_/k_2_, the coefficient of variation was significantly lower for K_1_/k_2_ and thus, it would be the preferred measure for quantification. There was also a discrepancy between the region-level and voxel-level results in [^15^O]H_2_O data in arteries, lungs, and the heart, which likely occurs due to high arterial volume fraction in these regions. Particularly, the K_1_ and V_A_ parameters of 1TCM are highly correlated in regions with high arterial volume fraction and cannot be reliably estimated. In such regions, it is possible to used fixed value (V_A_) for arterial volume fraction, which may help to overcome the parameter identifiability issues.

Model fits were generally satisfactory across regions, except in kidneys in [^15^O]H_2_O data, where slight underestimation was observed in the later time points, likely due to complex tracer kinetics behavior that is not described well by the standard [^15^O]H_2_O 1TCM.

Since this deviation occurs only in the later part of the scan, perfusion may still be reliably estimated using the earlier data. However, the pipeline is flexible and modular and tailored models with customized inputs can be included, but this warrants further research and careful validation.

## CONCLUSION

TurBO pipeline provides a reproducible, reliable and computationally fast approach for analyzing LAFOV PET data and enables interorgan interaction studies in tissue perfusion and metabolism.

## KEY POINTS

QUESTION:

How can LAFOV PET image processing and kinetic modelling be performed efficiently?

PERTINENT FINDINGS:

Based on the [^15^O]H_2_O (n=21) and [^18^F]FDG (n=16) validation data, the TurBO pipeline enables fast, accurate, and reproducible analysis of LAFOV PET perfusion and metabolism at both regional and voxel levels.

IMPLICATIONS FOR PATIENT CARE:

TurBO offers a robust and time-saving solution for processing and analyzing LAFOV PET data, which could be utilized also for the clinical data.

