## Supplemental material for "Automated Long Axial Field of View PET Image Processing and Kinetic Modelling with the TurBO Toolbox"

**SUPPLEMENTARY MATERIAL**

### MATERIALS AND METHODS

#### *LAFOV [$^{15}$O]$H_2O$ PET data modelling*

For [$^{15}$O]$H_2O$, a one-tissue compartmental model (1TCM) (*1*) is fitted for each measured regional time activity curve. The model is defined using the following equations (see subsection "[$^{15}$O]$H_2O$ model equations" for details):

$$C_\mathrm{T}(T) = K_1 \int_0^T C_\mathrm{A}(t) - k_2 \int_0^T C_\mathrm{T}(t)$$
$$C_\mathrm{PET}(t) = C_\mathrm{T}(t) + V_\mathrm{A} C_\mathrm{A}(t)$$

where $C_{PET}$ is the measured PET activity concentration, $C_T$ is the tissue activity concentration, $C_A$ is the arterial activity concentration corrected for radiotracer delay in tissue, $V_A$ is the arterial volume fraction, $K_1$=f describes the blood flow, and $k_2$=f/p, where p=$K_1/k_2$ is the partition coefficient of water (i.e., distribution volume).

Because the standard 1TCM model fitting with a descending aorta IDIF is not feasible for the liver, the arterial and portal vein blood flow fractions $r_a=f_a/(f_a+f_p)$ and $r_p=f_p/(f_a+f_p)$, where $f_a$ and $f_p$ represent arterial and portal vein blood flow (*2*), are first estimated using the dual-input model (*3*) with the descending aorta IDIF $C_{IDIF}$ and a portal vein input curve $C_{PV}$ (that is estimated using $C_{IDIF}$). Consequently, the resulting combined input

$$C_\mathrm{liverIF}(t) = r_\mathrm{a}\, C_\mathrm{IDIF}(t) + r_\mathrm{p}\, C_\mathrm{pv}(t)$$

is used for estimating liver parameters with the standard 1TCM.

Voxel-level $[^{15}O]H_2O$ quantification is carried out using non-negative least squares (NNLS, Lawson and Hanson, 1995) either by estimating all three 1TCM model parameters ($K_1$, $k_2$, $V_A$), or basis function approach (*5*), where discrete $k_2$ values are used for calculating the basis functions, and $K_1$, $V_A$ are estimated with NNLS (see subsection "$[^{15}O]H_2O$ model equations" for details). For both voxel-level methods, the radiotracer delay parameter is estimated separately for each voxel, except in the liver with combined input, where the input and the delay are obtained from the ROI-level quantification.

*$[^{15}O]H_2O$ model equations*

One-tissue compartmental model is described using the following differential equation:

$$\frac{dC_T(t)}{dt} = K_1 C_A(t) - k_2 C_T(t) \;\; (Eq\ 1).$$

Assuming that initial concentration in tissue compartment is zero ($C_T(0) = 0$), eq (1) can be integrated to provide the tissue concentration at time $T$:

$$C_T(T) = K_1 \int_0^T C_A(t)dt - k_2 \int_0^T C_T(t)dt \;\; (Eq\ 2).$$

The measured PET radioactivity concentration in tissue is contaminated by spillover from adjacent blood vessels $C_B$ and vascular volume $V_B$ inside the measured region $C_{PET}(t)$, which can be formulated as follows:

$$C_{PET}(t) = V_B C_B(t) + C_T(t) \;\; \text{(Eq 3)}.$$

Substituting $C_T(t)$ in eq (3) with eq (2) gives the following equation:

$$C_{PET}(t) = V_B C_B(t) + K_1 \int_0^T C_A(t)dt - k_2 \int_0^T C_T(t)dt \;\; (Eq\ 4).$$

Integration and rearrangement of eq (3) gives

$$\int_0^T C_T(t)dt = \int_0^T C_{PET}(t)dt - V_B \int_0^T C_B(t)dt \;\; (Eq\ 5),$$

which is then substituted in eq (4), providing the following equation:

$$C_{PET}(t) = V_B C_B(t) + K_1 \int_0^T C_A(t)dt - k_2 \int_0^T C_{PET}(t)dt + k_2 V_B \int_0^T C_B(t)dt \;\; (Eq\ 6).$$

If the radioactivity concentration in blood can be represented by the model input function, as is the case with $[^{15}O]H_2O$, then $C_B(t) = C_A(t)$ and $V_B = V_A$, and then eq (6) simplifies to eq (7):

$$C_{PET}(t) = V_A C_A(t) + (K_1 + V_A k_2) \int_0^T C_A(t)dt - k_2 \int_0^T C_{PET}(t)dt \;\; \text{(Eq 7)}.$$

Equation (7) is a multilinear equation $y = p_1\, x_1 + \; p_2 x_2 + \; p_3 x_3$, where the coefficients

$$V_A = p_1$$
$$k_2 = p_3$$
$$K_1 = p_2 - V_A k_2$$

can be estimated with least squares method (*6*). Particularly, non-negative least squares (NNLS, Lawson and Hanson, 1995) is well-suited method for estimating the coefficients.

The differential equation (1) can also be directly solved using convolution:

$$C_T(t) = (1 - V_A) K_1 C_A * e^{-k_2 t} \; (Eq\ 8),$$

where $C_T(t)$ is the radioactivity concentration in tissue, $C_A(t)$ is the arterial input function and * is the convolution operator. By denoting $C_B(t) = C_A(t)$ and $V_B = V_A$, and substituting eq (8) to eq (3), we get

$$C_{PET}(t) = (1 - V_A)K_1 C_A * e^{-k_2 t} + V_A C_A(t) \ (Eq\ 9).$$

The equation (9) can be rewritten as a multilinear equation (*7*):

$$C_T(t) = \theta_1 B(k_2, t) + \theta_2 C_A(t),$$

using basis functions

$$B(k_2, t) = C_A(t) * e^{-k_2 t},$$

and where the coefficients

$$\theta_1 = (1 - V_A)K_1$$
$$\theta_2 = V_A$$

can be estimated with NNLS.

*LAFOV [$^{18}$F]FDG PET data modelling*

To quantify glucose metabolism using [$^{18}$F]FDG, the descending aorta IDIF is first converted to plasma (*8*) using the individual hematocrit measurement, after which regional and voxel-level Patlak plot (*9*), fractional uptake ratio (FUR) (*10*) and standardised uptake value (SUV) estimates can be calculated similarly in all regions. Because lumped constant (LC) values that are used in conversion from [$^{18}$F]FDG to glucose uptake is still unknown for most tissue types, we used published LC values only for specific organs (*11*) ($LC_{brain}$=0.65, $LC_{myocardium}$=1, $LC_{muscles}$=1.16), and LC=1 for all other tissues.

*Data acquisition*

The subject's demographic details are listed in **Table S1**.

**Table S1.** Study sample characteristics.

| Demographic | [$^{15}$O]$H_2O$ | [$^{18}$F]FDG |
|---|---|---|
| n (Males/Females) | 8 / 13 | 4 / 12 |
| Age y (mean ± sd) | 65.4 ± 8.9 | 38.0 ± 7.0 |
| Dose MBq (mean ± sd) | 359.6 ± 23.8 | 171.8 ± 8.8 |
| BMI kg/m2 (mean ± sd) | 28.0 ± 4.9 | 30.6 ± 9.2 |

[$^{15}$O]$H_2O$ PET data were acquired for 280 s following 359.6 ± 23.8 MBq bolus injection over 10–15 seconds (Radiowater Generator, Hidex Oy, Finland). Imaging started 30 seconds after the start of the bolus injection. The data were reconstructed into 24 frames (14 × 5 s, 3 × 10 s, 3 × 20 s, 4 × 30 s) using an image matrix of 220 × 220 × 380 and a voxel size of 1.65 × 1.65 × 2.80 $mm^3$. Reconstruction was performed with an ordered-subsets expectation maximization (OSEM) algorithm (3 iterations, 5 subsets) using point-spread function and time-of-flight modelling, and included corrections for decay, randoms, attenuation, and scatter.

[$^{18}$F]FDG PET data were acquired for 55 minutes during hyperinsulinemic, euglycemic clamp after 171.8 ± 8.8 MBq bolus injection. Before the scan, two venous catheters were inserted in the opposite forearms - one for the insulin and glucose infusions and for injecting

[$^{18}$F]FDG, and the other for collecting venous blood samples, arterialised by placing a hot water bottle distally on the arm. After the collection of fasting plasma blood samples, hyperinsulinemic, euglycemic clamp was started (*12*). Insulin (Actrapid, Novo Nordisk A/S, Bagsvaerd, Denmark) was administered with a dose of 40 mU/min/m$^2$ of body surface area, and a variable rate of 20% glucose was infused based on plasma glucose measurements performed every 5–10 min to maintain euglycemia (plasma glucose 5.0 mmol/L). The [$^{18}$F]FDG PET scan was started 60 minutes after the start of insulin infusion under steady euglycemia. Data were reconstructed using OSEM algorithm (4 iterations, 5 subsets) using point-spread function and time-of-flight modelling, and included corrections for decay, randoms, attenuation, and scatter. The data were reconstructed into 34 frames (l2 x 5s, 6 x10s, 6 x 20s, 2 x 60s, 2 x 120s, 4 x 300s, 2x 600s) with 440 x 440 x 354 matrix size and 1.65 x 1.65 x 3.0 mm$^3$ voxel-size.

Prior to the PET image acquisition, the total-body CT images were acquired and reconstructed to 512 × 512 × 380 image matrix with a voxel size of 0.977 × 0.977 × 2.80 mm$^3$.

*Quality control plots for representative [$^{15}$O]$H_2O$ subject:*

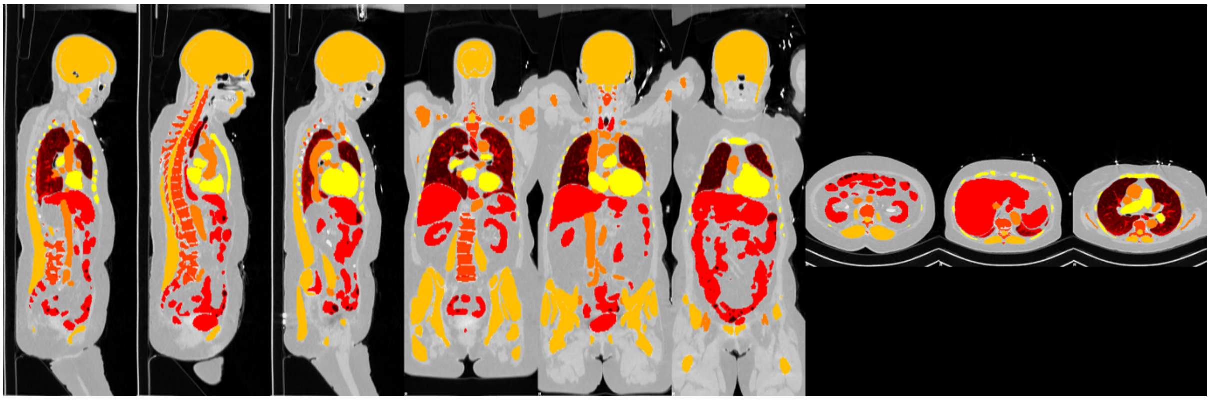

**Figure S1.** CT segments overlaid on CT image.

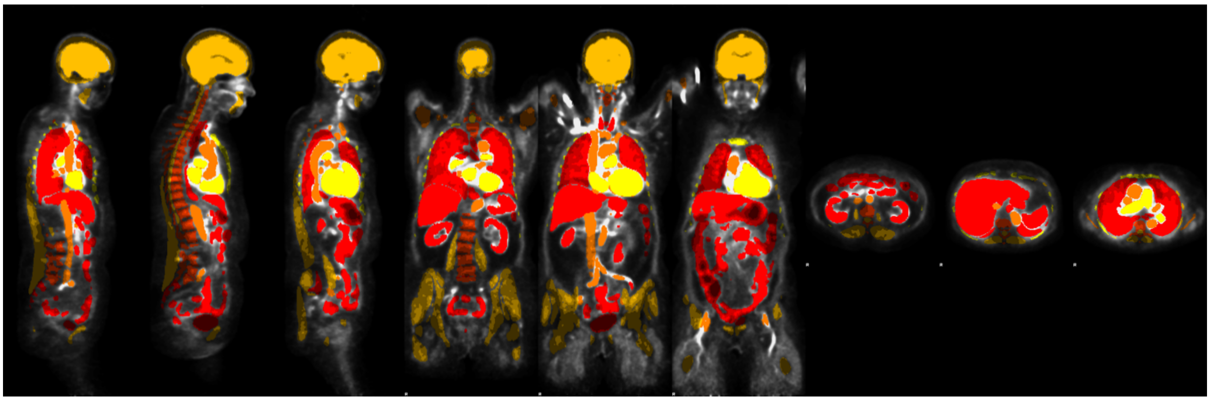

**Figure S2.** CT segments overlaid on mean PET image.

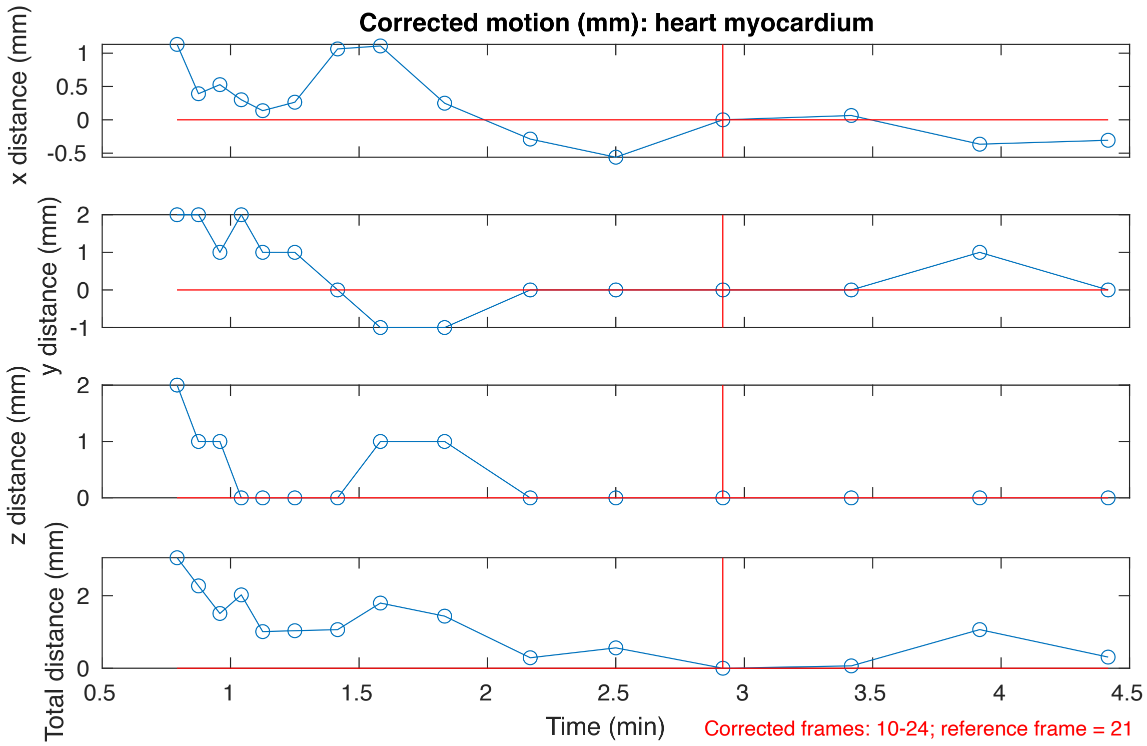


**Figure S3.** Corrected motion for heart myocardium

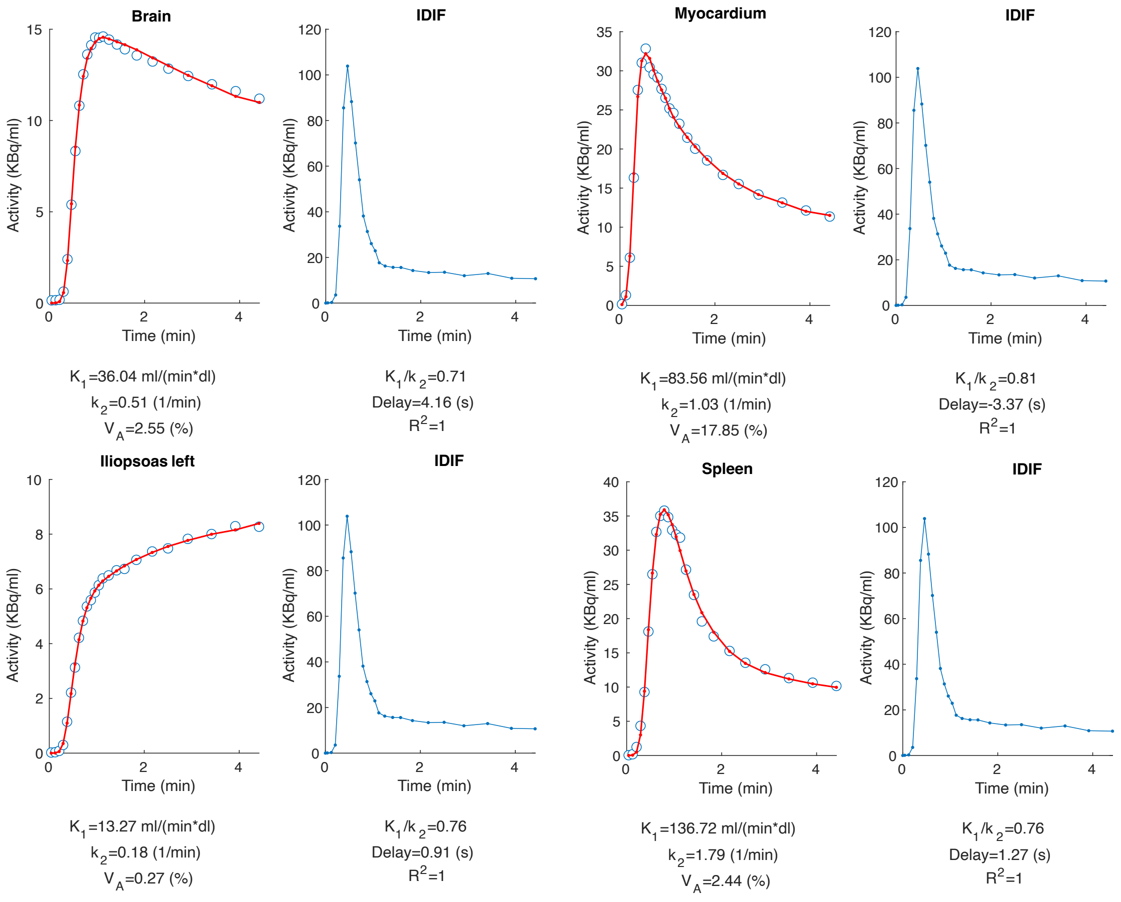


**Figure S4.** Example model fits

### RESULTS

**Table S2.** Typical TURBO-pipeline processing times time for representative cases of [$^{15}$O]$H_2O$ and [$^{18}$F]FDG data. Average processing times were measured using data from 3 subjects on our computational server using 5 parallel processes (details below), with CT segmentation done on a GPU. The processing time depends on the number of frames (24 for [$^{15}$O]$H_2O$ and 34 for [$^{18}$F]FDG), image matrix size (220 × 220 × 380 for [$^{15}$O]$H_2O$ and 440 x 440 x 354 for [$^{18}$F]FDG) and the available computational resources. The TURBO pipeline supports parallelization either by the number of image frames, or by the number of subjects.

| **Process** | **[$^{15}$O]$H_2O$ processing time (min)** | **[$^{18}$F]FDG processing time (min)** |
|---|---|---|
| CT segmentation | 5 | 7 |
| CT coregistration | 6 | 17 |
| PET motion correction | 17 | 40 |
| TAC extraction | 1 | 7 |
| Input data processing | 1 | 2 |
| Quality control | 2 | 3 |
| ROI-level modelling | 2 | 0.5 |
| Voxel-level modelling | 20 | 3 |
| Magia brain processing & modelling | 7 | 6 |
| Total running time | 61 | 85.5 |

System: Intel(R) Xeon(R) Platinum 8468V (97.5M Cache, 2.40 GHz, 879 Gb RAM) with NVIDIA L40 GPU (46 GB VRAM).

**Table S3.** Input ROI volumes.

| | **[$^{15}$O]$H_2O$** | | | **[$^{18}$F]FDG** | | |
|---|---|---|---|---|---|---|
| **Measure** | **Manual input** | **IDIF** | **p-value*** | **Manual input** | **IDIF** | **p-value*** |
| Volume (ml) | 4.0 (2.2) | 4.0 (0.7) | 0.99 | 8.2 (2.0) | 4.5 (0.9) | p < 0.01 |
| AUC/1000 | 120.7 (28.2) | 120.4 (28.8) | 0.53 | 167.5 (29.2) | 167.3 (30.1) | 0.84 |
| Dice coefficient | 0.5 (0.1) | | | 0.5 (0.1) | | |

*paired t-test

**Table S4.** Corrected frame-wise motion (mm) for selected six regions in [$^{15}$O]$H_2O$ and [$^{18}$F]FDG test data.

| Region | [$^{15}$O]$H_2O$ | | [$^{18}$F]FDG | |
|---|---|---|---|---|
| | Mean motion (mm) mean (sd) | Max motion (mm) mean (sd) | Mean motion (mm) mean (sd) | Max motion (mm) mean (sd) |
| Brain GMctx | 0.6 (0.6) | 2.1 (1.0) | 2.3 (0.9) | 6.1 (2.0) |
| Left iliopsoas | 2.4 (0.7) | 4.7 (1.2) | 3.8 (1.9) | 6.9 (3.3) |
| Liver | 5.0 (1.9) | 18.7 (7.8) | 4.3 (1.9) | 18.5 (17.6) |
| Right kidney | 5.1 (1.5) | 16.4 (7.5) | 4.1 (1.5) | 9.9 (3.1) |
| Spleen | 3.1 (0.8) | 6.5 (3.3) | 3.9 (1.2) | 8.0 (2.6 |
| Pancreas | 4.4 (1.4) | 11.1 (6.1) | 4.6 (1.4) | 13.5 (6.1) |

**Table S5.** Volumes of manually drawn ROIs and segmented CT ROIs.

| Radioligand | Region | Segmented CT volume (ml) mean (sd) | Manual ROI volume (ml) mean (sd) |
|---|---|---|---|
| [$^{15}$O]$H_2O$ | Liver | 1510.7 (464.6) | 42.4 (20.6) |
| | Spleen | 181.2 (63.7) | 13.9 (5.3) |
| | Kidney | 382.6 (137.0) | 47.4 (16.9) |
| [$^{18}$F]FDG | Liver | 1639.8 (330.5) | 29.6 (12.9) |
| | Iliopsoas | 578.9 (92.3) | 11.6 (4.5) |

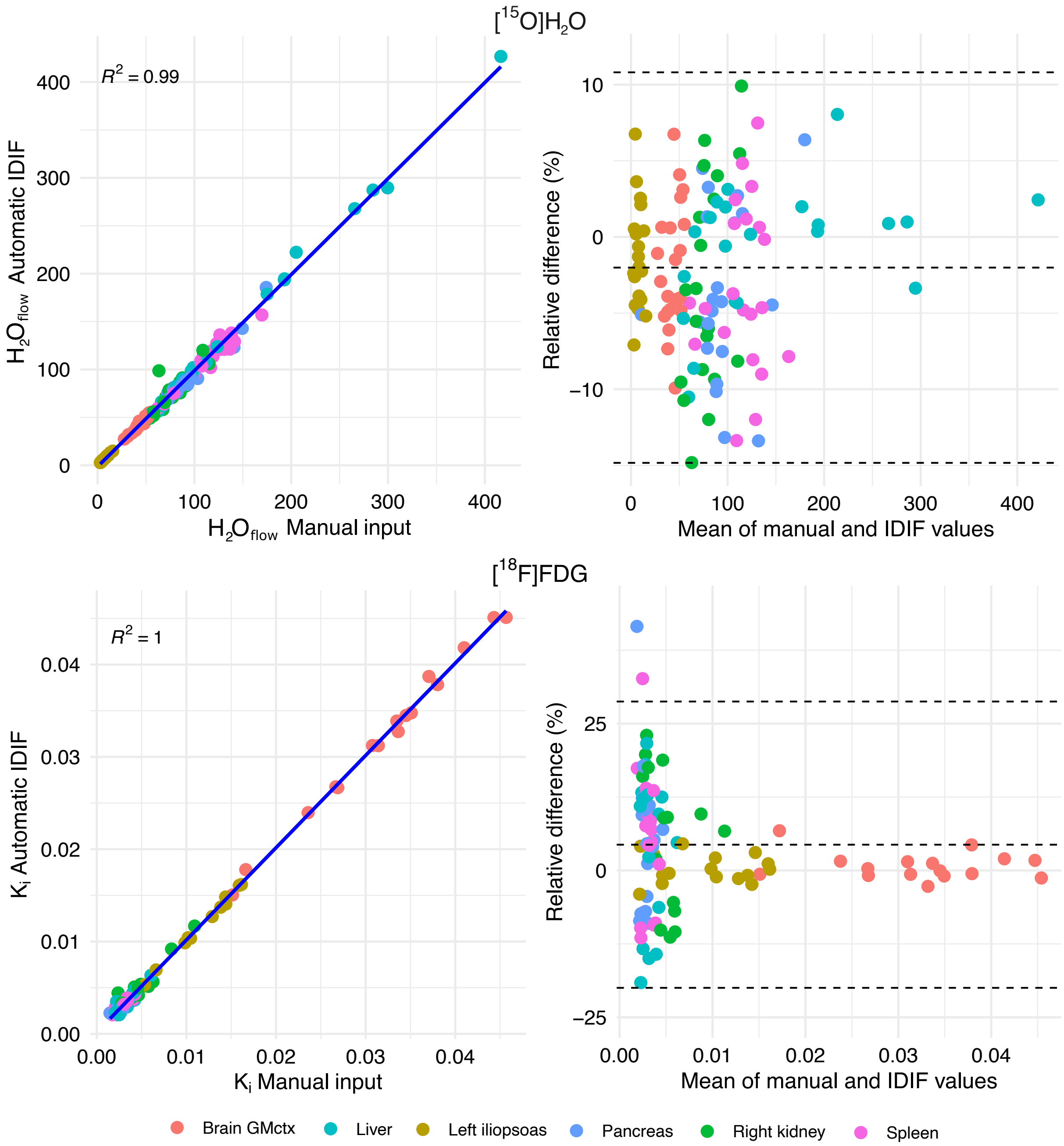


**Figure S4.** Pearson's correlation and Bland-Altman plots of parameter estimates ($H_2O_{flow}$ = ($K_1(1-V_A)$) for [$^{15}$O]$H_2O$ and Patlak $K_i$ for [$^{18}$F]FDG) in six example regions, calculated using manually derived, and image derived input (IDIF) from descending aorta.

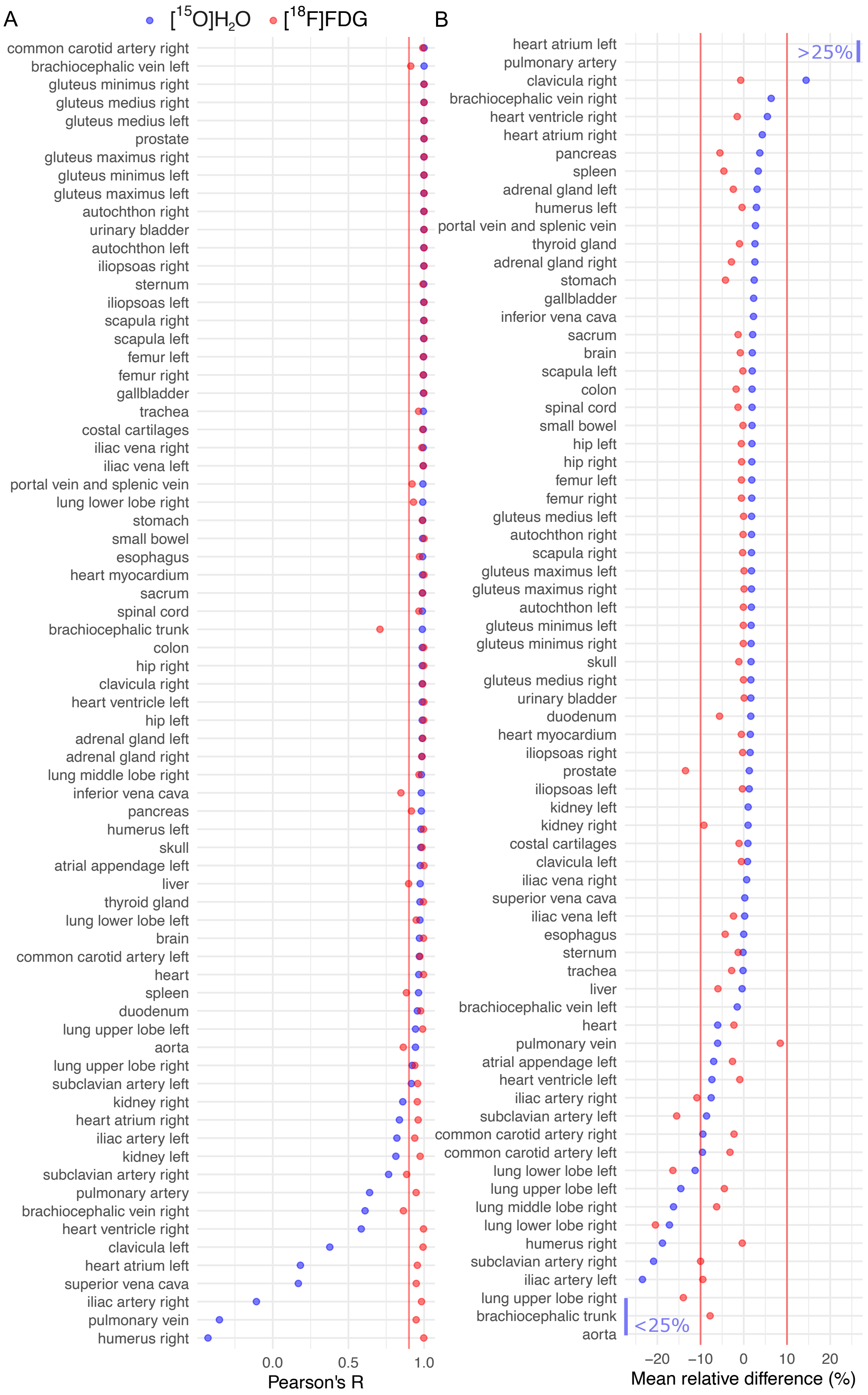


**Figure S5. A)** Pearson's correlation of model parameter estimates calculated using manually derived, and image derived input (IDIF) from descending aorta. **B)** Mean relative differences between model parameter estimates calculated using manually derived, and image derived input (IDIF) from descending aorta. Regional $H_2O_{flow} = K_1(1-V_A)$ estimates for $[^{15}O]H_2O$ are coloured in blue and $[^{18}F]FDG$ $K_i$ estimates in red.

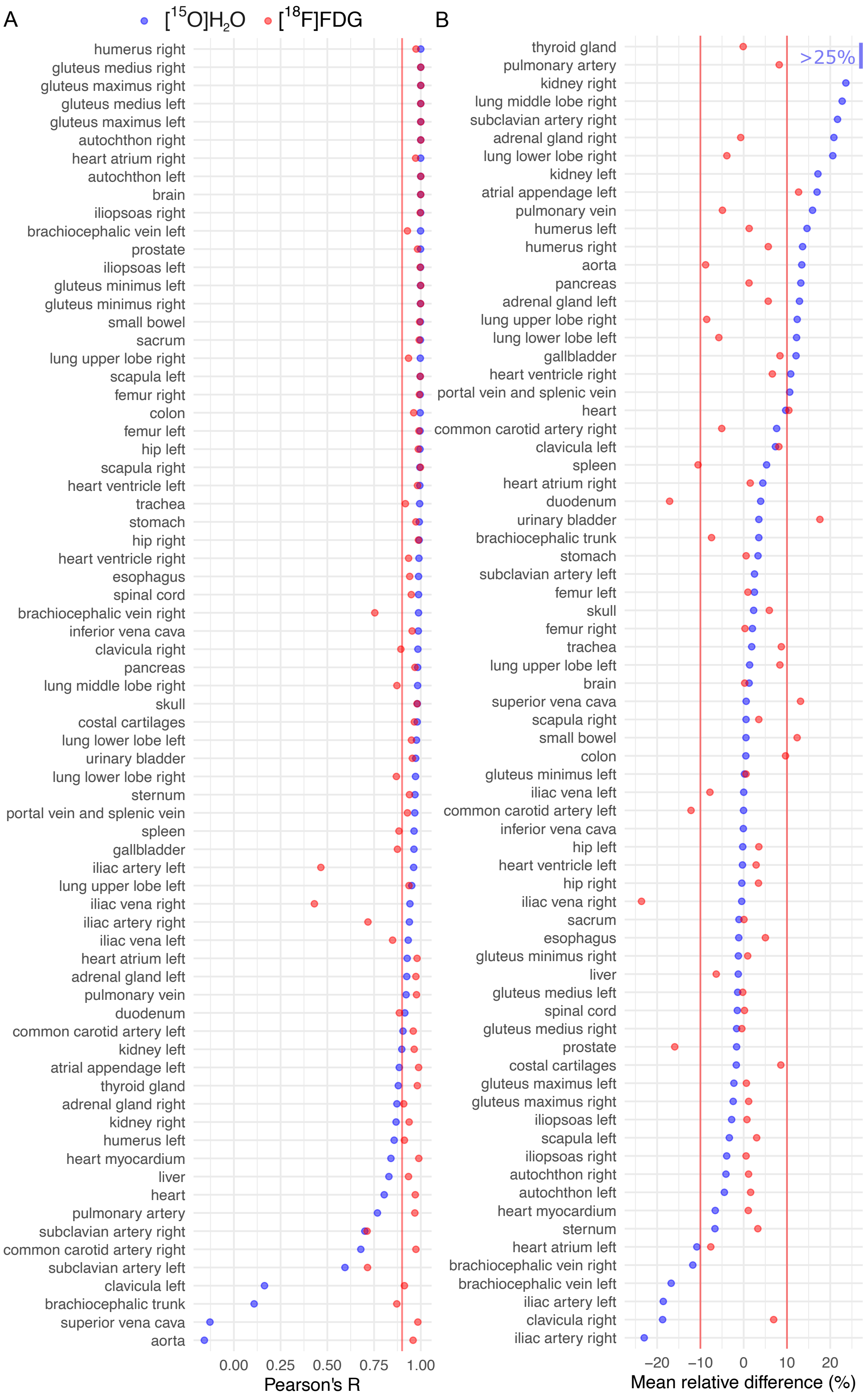


**Figure S6. A)** Pearson's correlation of model parameter estimates calculated using data with and without motion correction. **B)** Mean relative differences between model parameter estimates calculated using data with and without motion correction. Regional $H_2O_{flow} = K_1(1-V_A)$ estimates for $[^{15}O]H_2O$ are coloured in blue and $[^{18}F]$FDG $K_i$ estimates in red.

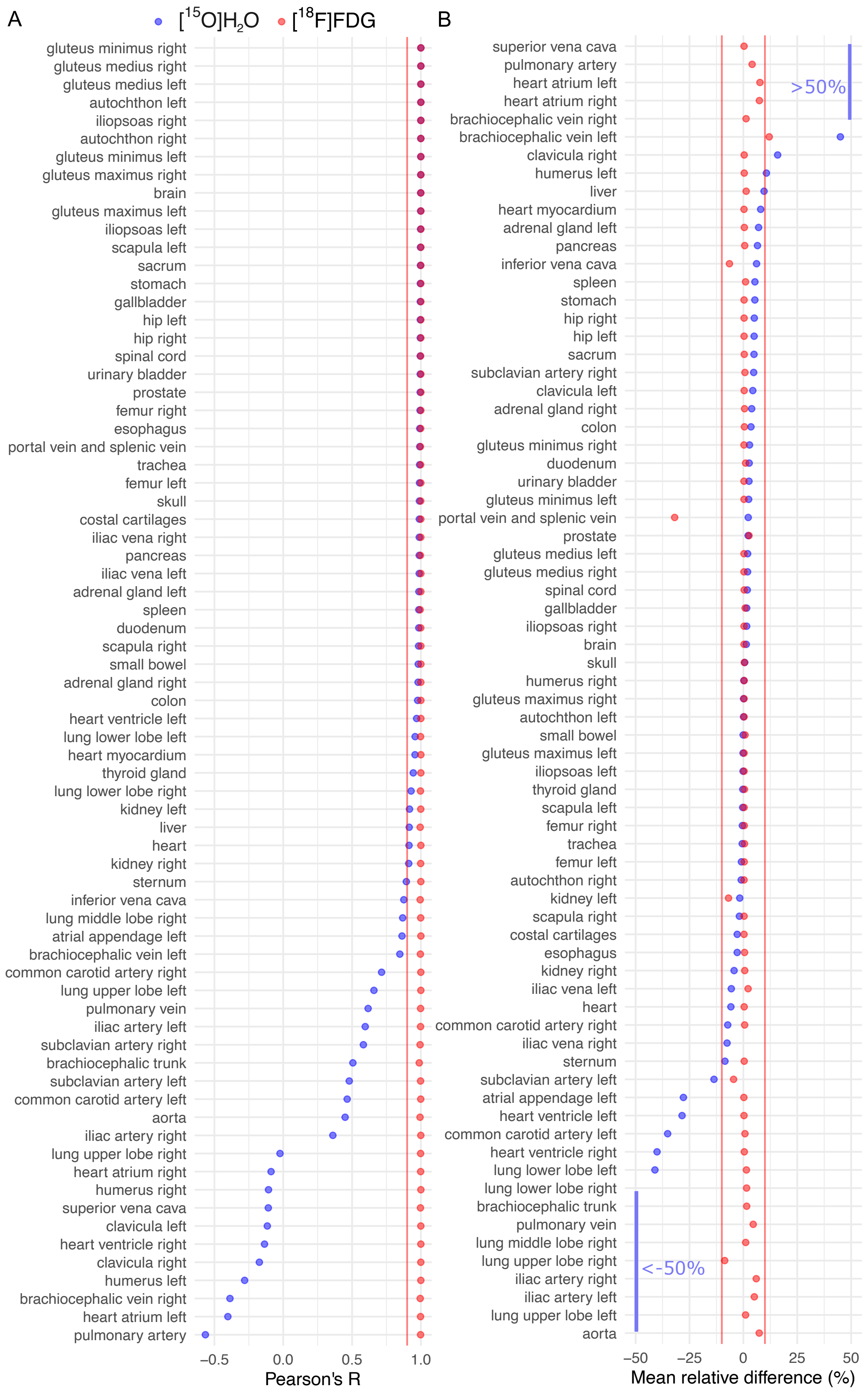


**Figure S7. A)** Pearson's correlation of model parameter estimates calculated using ROI- and voxel level modelling. Basis function method was used at voxel-level. **B)** Mean relative differences between model parameter estimates calculated using ROI- and voxel level modelling**.** Regional $H_2O_{flow} = K_1(1-V_A)$ estimates for $[^{15}O]H_2O$ are coloured in blue and $[^{18}F]FDG$ $K_i$ estimates in red.

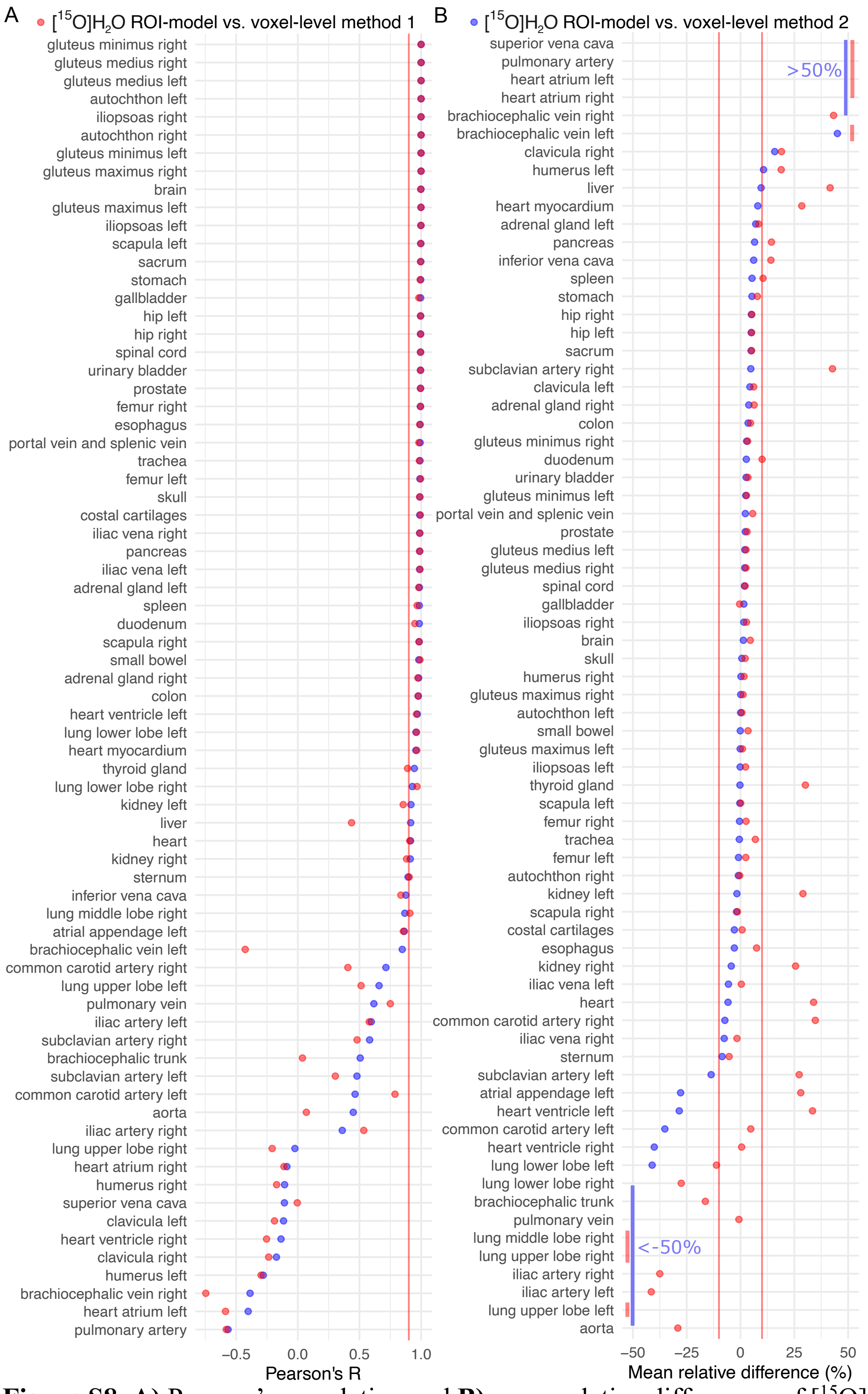


**Figure S8. A)** Pearson's correlation and **B)** mean relative differences of [15O]$H_2O$ 1-tissue compartment model $H_2O_{flow} = K_1(1-V_A)$ estimates calculated using ROI- and voxel level modelling. Red markers show the comparison between a three-parameter ($K_1$, $k_2$, $V_A$) ROI-level model and three-parameter voxel-level model (method 1), and blue markers show the comparison between three-parameter ROI level-model and basis function voxel-level model (method 2) with two estimated model parameters ($K_1$, $V_A$).

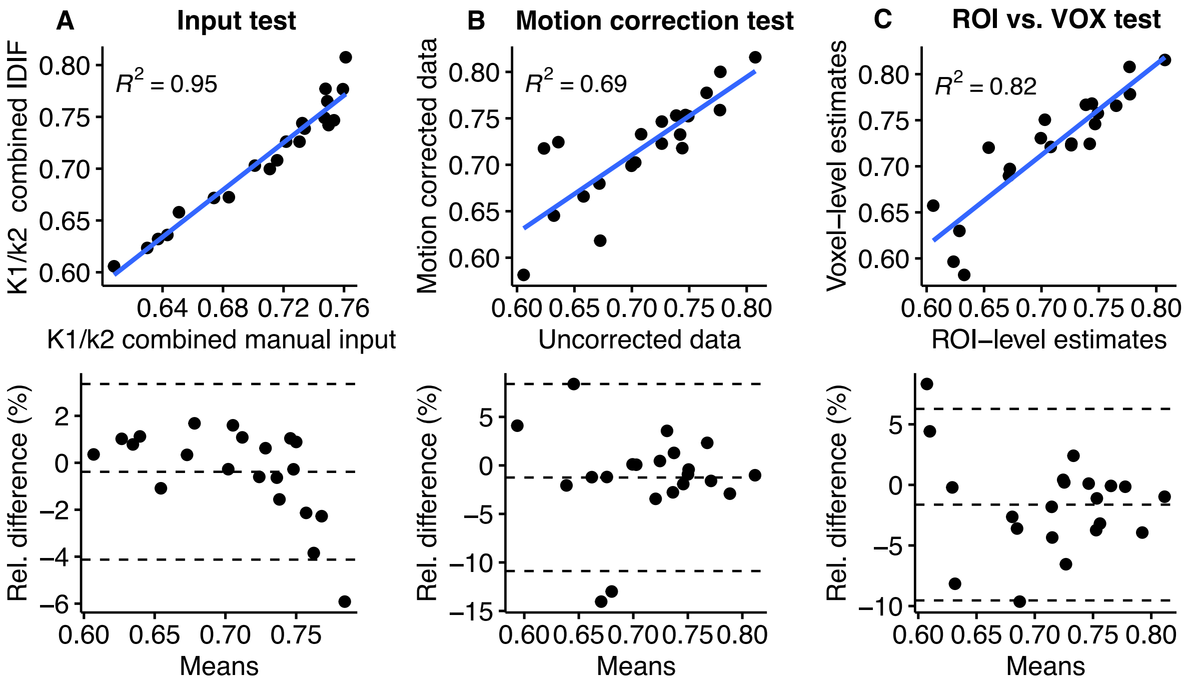


**Figure S9.** Scatterplots and Bland-Altman plots illustrating correlation and relative difference between $[^{15}O]H_2O$ liver partition coefficient of water ($p=K_1/k_2$) estimates, where **A**) parameter estimates obtained with manual input are compared with parameter estimates calculated using IDIF, **B**) parameter estimates calculated using uncorrected data are compared with parameter estimates calculated using motion corrected data, and **C**) ROI-level parameter estimates are compared with voxel-level parameter estimates.
